# Switching between bacteriostatic and bactericidal antimicrobials for retreatment of bovine respiratory disease (BRD) relapses is associated with an increased frequency of resistant pathogen isolation from veterinary diagnostic laboratory submissions

**DOI:** 10.1101/675066

**Authors:** Johann F. Coetzee, Drew R. Magstadt, Lendie Follett, Pritam K. Sidhu, Adlai M. Schuler, Adam C. Krull, Vickie L. Cooper, Terry J. Engelken, Annette M. O’Connor

**Affiliations:** Department of Veterinary Diagnostic and Production Animal Medicine, College of Veterinary Medicine, Iowa State University, Ames, IA, 50011; Department of Information Management and Business Analytics, College of Business and Public Administration, Drake University, Des Moines, IA, 50311; Department of Anatomy and Physiology, College of Veterinary Medicine, Kansas State University, Manhattan, KS, 66506

**Keywords:** antimicrobials, bacteriostatic, bactericidal, bovine respiratory disease, *Histophilus somni*, *Mannheimia haemolytica*, *Pasteurella multocida*, resistance.

## Abstract

Although 90% of BRD relapses are reported to receive retreatment with a different class of antimicrobial, studies examining the impact of antimicrobial selection (i.e. bactericidal or bacteriostatic) on retreatment outcomes and the emergence of antimicrobial resistance (AMR) are deficient in the published literature. A survey was conducted to determine the association between antimicrobial class selection for retreatment of BRD relapses on antimicrobial susceptibility of *Mannheimia haemolytica*, *Pasteurella multocida*, and *Histophilus somni.* Pathogens were isolated from samples submitted to the Iowa State University Veterinary Diagnostic Laboratory from January 2013 to December 2015. A total of 781 isolates with corresponding animal case histories, including treatment protocols, were included in the analysis. Original susceptibility testing of these isolates for ceftiofur, danofloxacin, enrofloxacin, florfenicol, oxytetracycline, spectinomycin, tilmicosin, and tulathromycin was performed using Clinical and Laboratory Standards Institute guidelines. Data were analyzed using a Bayesian approach to evaluate whether retreatment with antimicrobials of different mechanistic classes (bactericidal or bacteriostatic) increased the probability of resistant BRD pathogen isolation in calves. The posterior distribution we calculated suggests that an increased number of treatments is associated with a greater probability of isolates resistant to at least one antimicrobial. In addition, the frequency of resistant *M. haemolytica* isolates was greater with retreatment using antimicrobials of different mechanistic classes than retreatment with the same class. Specifically, treatment protocols using a bacteriostatic drug first followed by retreatment with a bactericidal drug was associated with a higher frequency of resistant BRD pathogen isolation. This effect was more profound with specific treatment combinations; tulathromycin (bacteriostatic) followed by ceftiofur (bactericidal) was associated with the highest probability of resistant isolates among all antimicrobial combinations. These findings suggest that the selection of antimicrobial mechanistic class for retreatment of BRD should be considered as part of an antimicrobial stewardship program.

## Introduction

Bovine respiratory disease (BRD) is one of the most important diseases facing the beef cattle industry [1]. Annual economic losses due to BRD are estimated to approach $1 billion in the United States alone [1, 2]. Treatment and control of BRD are currently predicated on administration of antimicrobial therapy directed toward the primary bacterial pathogens *Mannheimia haemolytica*, *Pasteurella multocida*, and *Histophilus somni*. Antimicrobial drugs are broadly classified into two groups, namely those that inhibit growth of the organism (ie. bacteriostatic) and those that kill the organism (ie, bactericidal). The National Animal Health Monitoring System Feedlot 2011 study reported that 21.2 ± 2.0% (standard error; SE) of cattle in feedlots were administered antimicrobials to control an expected outbreak of BRD, and approximately 15% of feedlot cattle required a second antimicrobial treatment for the disease [3,4,5]. Although approximately 90% of cases with BRD relapse were reported to receive retreatment with a different antimicrobial mechanistic class [5], studies examining the impact of antimicrobial drug class on retreatment outcomes and the emergence of antimicrobial resistance (AMR) are scarce in the published literature. Knowledge of the impact of antimicrobial drug selection on AMR emergence is needed to develop judicious use guidelines that preserve antimicrobial efficacy and advance antimicrobial stewardship.

Minimum inhibitory concentration (MIC) data obtained from samples submitted to veterinary diagnostic laboratories (VDLs) reflect antimicrobial susceptibility and are commonly used to describe AMR changes in livestock populations [6,7,8]. A retrospective study of *M. haemolytica,* recovered from lung samples submitted to the Kansas State University VDL between 2009 and 2011, reported a 7-fold increase in the number of isolates resistant to five or more antimicrobials over a 3-year period [9]. However, the association between antimicrobial treatment and the recovery of a resistant *M. haemolytica* isolate could not be evaluated because individual animal treatment histories were not reported. Recently, our group reported an association between treatment history and antimicrobial sensitivity results from bacterial isolates obtained from BRD cases submitted to the Iowa State University VDL (ISU-VDL) from 2013– 2015 [10]. Bacterial isolates from cattle that received antimicrobial treatment showed a higher incidence of antimicrobial resistance than isolates from untreated cattle. Furthermore, the percentage of resistant isolates increased with the number of antimicrobial treatments. However, the relationships between the antimicrobial drug class selected for initial treatment and retreatment as well as the frequency of AMR pathogen isolation were not investigated.

It was revealed more than 50 years ago that an overall reduction in antimicrobial efficacy occurs when antimicrobials that cause target organism death (i.e., bactericidal agents) are used in combination with antimicrobials that only inhibit bacterial replication (i.e., bacteriostatic agents) [11, 12]. The resulting drug antagonism is associated with poorer clinical outcomes [12–14]. These findings suggest that the choice of antimicrobial drug class (i.e., bactericidal or bacteriostatic) in cases of relapse and retreatment may be a critical control point for mitigating AMR in beef production systems. The objectives of this study were to use a Bayesian approach to 1) obtain the posterior distribution of the resistance patterns for the number of treatments (1, 2, 3, or 4+) administered to cases submitted to the ISU-VDL, and 2) test the hypothesis that antimicrobial resistant BRD pathogens are recovered more frequently from calves that received second-line treatment from a different antimicrobial class than from calves that received second-line treatment from the same antimicrobial class.

## Materials and Methods

### Study design

This cross-sectional study used data collected from the electronic and paper laboratory records of the ISU-VDL from January 1, 2013 to December 2, 2015, including the original documents, which were used to extract the relevant antimicrobial treatment information. The data were retrieved in 2016.

### Settings

The 1,251 isolates available for analysis were submitted to the ISU-VDL by referring veterinarians from 24 states. The majority of isolates were from Iowa (778), Minnesota (80), and South Dakota (49). Most isolates were obtained from animals housed in feedlots (498), confinement operations (268), or pastures (162). The demographic information from the sample submissions is summarized in Tables S1 and S2.

### Cases and case isolates

Bacterial isolate data and the corresponding case history information were included in the study upon meeting the following criteria: 1) The submitted samples were from a bovine field case (research cases were excluded); 2) *M. haemolytica*, *P. multocida*, or *H. somni* were isolated via routine culture; 3) The sample that yielded the isolate was from the lower respiratory tract (lung, pleural surface, bronchoalveolar lavage fluid); 4) MIC testing results were available; 5) The submission form stated a history of respiratory disease and/or evidence of pneumonia was described in autopsy findings or upon histological evaluation of lung tissue; and 6) The submitting veterinarian provided a treatment history that included either the generic or trade name of the antimicrobials used in the treatment of the case prior to sample submission.

### Study size

No *a priori* sample size was determined because the study was intended to be cross-sectional and hypothesis-generating. Therefore, sample size was determined solely by the number of eligible isolates available during the study period.

### Variables and data sources

The outcome of interest was the Clinical and Laboratory Standards Institute (CLSI)-validated interpretive category based on MIC.

Susceptibility testing was performed according to standard laboratory methods based on CLSI recommendations [15]. Briefly, the selected culture was grown overnight and a broth dilution was inoculated on a standard 96-well susceptibility plate (BOPO6F, Thermo Scientific, Oakwood Village, OH, USA) using an automated inoculation system (Sensititre AIM, Thermo Scientific). Susceptibility plates were read using a manual system (Sensititre Vizion System, Thermo Scientific) following 18–24 h incubation at 37°C.

Not all antimicrobial compounds included on the standard susceptibility plate have CLSI-validated interpretive breakpoints; therefore, only antimicrobials with CLSI-approved breakpoints [2] for respiratory disease caused by *M. haemolytica* were included in this study (S3 Table). The antimicrobials included in this study were ceftiofur, danofloxacin, enrofloxacin, florfenicol, oxytetracycline, spectinomycin, tilmicosin, and tulathromycin. Established CLSI-validated breakpoints are not available for tilmicosin against *P. multocida* or tilmicosin and danofloxacin against *H. somni* in BRD; therefore, these antimicrobials were included in this study using the CLSI-validated breakpoints for *M. haemolytica*.

Treatment history was recorded in the paper submission forms by the referring veterinarian. Information regarding the number of antimicrobial treatments, specific antimicrobials used, and non-antimicrobial treatments was manually extracted from these records by one investigator (AS). Isolates from submissions that explicitly stated no usage of antimicrobial drugs were assigned the treatment history classification of none (“0”). Isolates from cases in which information regarding antimicrobial treatments was unclear (e.g., “many” or “everything”) or not given were classified as “unknown.” Isolates from cases with treatment histories indicating the use of four or more antimicrobials were classified as “4+.”

Trade names were converted to generic drug names to determine the antimicrobial drug class (bacteriostatic or bactericidal) and the sequence of class administration for first- and second-line treatments. Drug class was assigned based on the established *in vitro* pharmacodynamics of the antimicrobial agent as summarized in Table 1.

Data on potential confounders or effect modifiers extracted from the submission form, including breed, sex, facility type, clinical signs, necropsy findings, vaccination status, and weights, were recorded (**Tables S1 and S2**). Finalized case report information, such as microscopic evidence of pneumonia, also was noted. Case information was classified as “unknown” if the information was not supplied or unclear. After each eligible record was identified, the submission forms for each case were individually reviewed by a single researcher (AS). Antimicrobial treatments were grouped as -cidal or -static based on antimicrobial activity level.

**Table 1.** Classification of antimicrobial drugs on the basis of antimicrobial activity.

| Bactericidal | Bacteriostatic |
| --- | --- |
| Ceftiofur | Chlortetracycline |
| Danofloxacin | Florfenicol |
| Enrofloxacin | Gamithromycin |
| Penicillin | Oxytetracycline |
|  | Spectinomycin |
|  | Sulfadimethoxine |
|  | Tildipirosin |
|  | Tilmicosin |
|  | Tulathromycin |
|  | Tylosin |

### Variable transformations

Due to sparse data for cases receiving multiple treatments, we arbitrarily chose to group together animals that received more than three treatments (4+). Animals with unknown treatment histories were excluded from the analysis.

For the subset of animals receiving just two treatments, we created two categorical variables. One categorical variable grouped the data into two levels: “same” to designate animals that received first- and second-line treatment from the same drug class (i.e., either bacteriostatic and bacteriostatic or bactericidal and bactericidal) and “different” to designate animals that received first- and second-line treatment from different drug classes (i.e., either bacteriostatic followed by bactericidal or bactericidal followed by bacteriostatic). We also created a four-level categorical variable to capture all possible combinations (4 levels: bacteriostatic followed by bactericidal, bacteriostatic followed by bacteriostatic, bactericidal followed by bacteriostatic, and bactericidal followed by bactericidal).

### Statistical analysis

Initial analysis included descriptive statistics to illustrate the distribution of the number of treatments cross tabulated with the number of antimicrobials of which the isolate was classified as resistant. The number of missing values also was determined. We created heat maps to show the pairwise interactions of antimicrobial treatment combinations associated with the development of resistant *M. haemolytica* mutants.

The approach for addressing our two objectives was to conduct a Bayesian analysis using a finite mixture model based on a zero-inflated beta-binomial model. The open source software R was used to conduct this analysis. For both objectives, we let *y_ij_* represent the number of resistant organisms, where *i* represents the level of the explanatory variable treatment and *j* represents the organism. We assume the observations are conditionally independent. We write the model as follows:

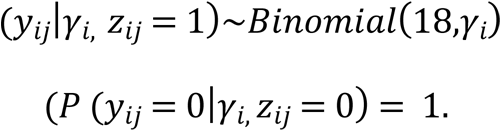

where *z_ij_* represents the category (i.e., antimicrobial drug class) and n = 18 represents the number of possible antimicrobials. Thus, the probability density function of *y_ij_* is:

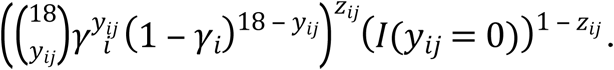

We allow the category indicator, *z_ij_*, to also be conditionally independent with the following distributional assumption:

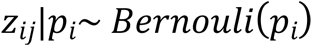

*Rho* (*p_i_*) and *gamma* (*γ_i_*) are assumed to be independent with priors specified as followed:

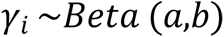

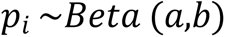

where *a* = 1 and *b* = 1.

For Objective 1, *i* in the model referred to the number of treatments reported by the submitter (i.e., *i* had five levels: 0, 1, 2, 3, and 4+ treatments). For Objective 2, two models were created for the subset of animals that received two treatments. For the first model (Objective 2 Model 1), γ_i_ referred to the two-level sequence of treatments reported by the submitter (i.e., *i* had two levels: same and different). For the second model (Objective 2 Model 2), γ_i_ referred to the four-level sequence of treatments reported by the submitter (i.e., *i* had four levels: bactericidal-bactericidal, bactericidal-bacteriostatic, bacteriostatic-bactericidal, and bacteriostatic-bacteriostatic). We sampled from the joint posterior distribution of γ_i_ and γ_i_ implied by the model using a Metropolis random walk Markov chain Monte Carlo (MCMC) approach.

One output from each model was the posterior distribution of ρ_i_ based on each *i*^th^ level of the explanatory variable; i.e., ρ_i_ is the probability that an animal in group *i* comes from the binomial distribution. We use this posterior distribution to make inferences about the data. For example, we are interested in the probability that an organism is resistant to at least one antimicrobial, which is given by ρ_i_* (probability binomial [18, γ_i_] random variable > 0). We are also interested in whether this probability is associated with either the number of times an animal is treated or the treatment sequence. The other output was the posterior distribution of γ_i_ based on each *i^th^* level of the explanatory variable. Posterior γ distributions that are shifted to the right indicate an isolate that is resistant to a higher number of antimicrobials.

To “test” the relationship between the probability of at least one resistant test result and the number of treatments, we determined the posterior distribution of *p_i+1_* > *p_i_*, i.e., how often the probability of having at least one resistant test was higher for *i + 1* treatments compared to *I* treatments.

After creating the models, we assessed several discrepancy measures, including the number of zeros in the posterior distribution of *y\** and the number of extreme values. This first reflects the inflation of zeros in the observed distribution and is mainly informed by the posterior distribution of *ρ_i._* The second measure assesses the distribution of resistant organisms, given that they are resistant, and is mainly informed by the posterior distribution of γ_i._ For each posterior distribution of γ_i_ and ρ_i_, we reported the 95% credible interval (95% CI) and also the posterior probability that γ_i <_ γ_≠i_ and ρ_i <_ρ_≠i._ for all possible pairwise comparisons.

### Ancillary analyses and sensitivity analyses

We originally intended to conduct further subgroup analysis based on the particular isolates of *M. haemolytica*, *P. multocida*, and *H. somni*. However, on further examination, we determined that the data were too sparse to warrant further subgrouping. We also were originally interested in the impact of two covariates, simultaneous viral and *Mycoplasma* spp. infection, on the posterior distribution; however, descriptive analysis indicated that this approach was unlikely to be useful. Thus, although we extracted these data, we did not conduct these analyses.

## Results and Discussion

In North America, BRD in feedlot cattle results in substantial economic losses due to the costs of treatment and deleterious effects on animal health and production [14, 16]. Although BRD has a complex, multifactorial etiology, *M. haemolytica*, *P. multocida*, and *H. somni* are most often associated with clinical disease [17, 18]. Therefore, use of antimicrobials is essential for the control and treatment of BRD in cattle. Commonly used antimicrobial agents that are approved in the US for treatment of BRD include ceftiofur, tilmicosin, tulathromycin, florfenicol, enrofloxacin, and danofloxacin.

An increasing number of reports regarding decreased efficacy of these antimicrobial agents for treatment of BRD have been published in recent years [9,19,20,21]. Typically, cattle affected with BRD are treated with a drug of a different antimicrobial class than the drug given for first treatment or disease prevention (metaphylaxis). When an animal does not respond to the first-line treatment, it may be treated with one or more additional classes of antimicrobial drugs over subsequent days. Data analyzed in the present study confirm the use of multiple classes of antimicrobial treatments used to treat BRD (Table 2), which is similar to the findings of an earlier report [10]. However, despite the frequent use of sequential treatments, there are no data indicating what drug classes, doses, or scheduling criteria might be optimal when using sequential treatments for BRD in cattle.

**Table 2.** Summary of bacterial isolates obtained from submitted samples of animals treated with bacteriostatic/bactericidal antimicrobial agents.

| Year | 2013 |  |  | 2014 |  |  | 2015 |  |  | Total |
| --- | --- | --- | --- | --- | --- | --- | --- | --- | --- | --- |
| Organisms (culture) | MH | PM | HS | MH | PM | HS | MH | PM | HS |  |
| Isolates from submissions with treatment history | 113 | 56 | 52 | 127 | 90 | 81 | 106 | 94 | 62 | 781 |
| Total isolates/year | 221 |  |  | 298 |  |  | 262 |  |  |  |
| Isolates from non-treated cases | 28 | 19 | 13 | 27 | 29 | 17 | 25 | 19 | 17 | 194 |
|  | 60 |  |  | 73 |  |  | 61 |  |  |  |
| Isolates from first-line bactericidal treatment |  |  |  |  |  |  |  |  |  |  |
| Ampicillin | 0 | 0 | 0 | 1 | 0 | 0 | 0 | 2 | 0 | 3 |
| Ceftiofur | 14 | 1 | 5 | 15 | 10 | 11 | 13 | 8 | 4 | 81 |
| Danofloxacin | 0 | 1 | 0 | 1 | 1 | 0 | 0 | 0 | 0 | 3 |
| Enrofloxacin | 12 | 5 | 6 | 10 | 9 | 11 | 13 | 14 | 5 | 85 |
| Penicillin | 2 | 2 | 0 | 0 | 0 | 0 | 2 | 2 | 1 | 9 |
| Total | 48 |  |  | 69 |  |  | 64 |  |  | 181 |
| Isolates from first-line bacteriostatic treatment |  |  |  |  |  |  |  |  |  |  |
| Chlortetracycline | 0 | 0 | 0 | 4 | 4 | 1 | 1 | 0 | 1 | 11 |
| Florfenicol | 6 | 6 | 7 | 8 | 7 | 9 | 16 | 8 | 8 | 75 |
| Gamithromycin | 2 | 1 | 0 | 9 | 6 | 2 | 5 | 1 | 2 | 28 |
| Oxytetracycline | 2 | 1 | 1 | 0 | 3 | 1 | 1 | 2 | 1 | 12 |
| Sulfadimethoxine | 2 | 2 | 0 | 0 | 0 | 1 | 1 | 0 | 0 | 6 |
| Tetracycline | 0 | 0 | 0 | 5 | 2 | 2 | 0 | 1 | 2 | 12 |
| Tildipirosin | 11 | 3 | 4 | 6 | 5 | 6 | 4 | 8 | 3 | 50 |
| Tilmicosin | 4 | 4 | 4 | 8 | 1 | 5 | 2 | 2 | 0 | 30 |
| Tulathromycin | 29 | 10 | 11 | 33 | 13 | 15 | 24 | 26 | 18 | 179 |
| Tylosin | 1 | 1 | 1 | 0 | 0 | 0 | 0 | 0 | 0 | 3 |
| Total | 113 |  |  | 156 |  |  | 137 |  |  | 406 |

(MH = *M. haemolytica*, PM = *P. multocida*, HS = *H. somni*)

A total of 1,251 bacterial isolates were available for our analysis, including 540 isolates of *M. haemolytica*, 404 isolates of *P. multocida*, and 307 isolates of *H. somni*. Isolates were obtained from 1,031 individual animals under 989 case submissions by 378 veterinarians. Table 2 summarizes the numbers of each organism isolated each year over the course of the study.

The full data set of 781 of 1,251 bacterial isolates was used for analysis because 470 isolates did not have treatment information included with the sample submission. The remaining dataset available for Objective 1 included 781 isolates, of which 194 received 0 treatments, 276 received 1 treatment, 211 received 2 treatments, 74 received 3 treatments, 23 received 4 treatments, 2 received 5 treatments, and 1 received 7 treatments. Missing data for this subset is presented in **Table S1**. Previous laboratory studies also identified multiple drug resistant (MDR) isolates from lung tissues collected from fatal BRD cases [9,14,22,23]. Poor response to antimicrobial therapy in fatal BRD cases may be associated with the presence of sub-inhibitory concentrations of antimicrobial drugs due to pre-treatment, which could induce positive selection leading to resistance [24, 25]. A total of 211 isolates were from animals that received only two treatments. Of these isolates, 101 were *M. haemolytica*, 50 were *H. somni*, and 60 were *P. multocida*. These isolates were treated with the same drug class in 97 cases (18 bactericidal-bactericidal and 79 bacteriostatic-bacteriostatic) and 114 were treated with different drug classes (52 bactericidal-bacteriostatic and 62 bacteriostatic-bactericidal).

The observed antimicrobial susceptibility profiles for *M. haemolytica* based on MIC data of cattle administered either the “same” (first and second treatment were both either bactericidal drugs, or bacteriostatic drugs) or “different” (first treatment was bactericidal and second was bacteriostatic or *vice versa*) antimicrobial treatment is presented in Fig. 1. A similar examination of the data was not conducted for *P. multocida* and *H. somni* because there were an insufficient number of isolates for this to be meaningful.

**Fig. 1.**
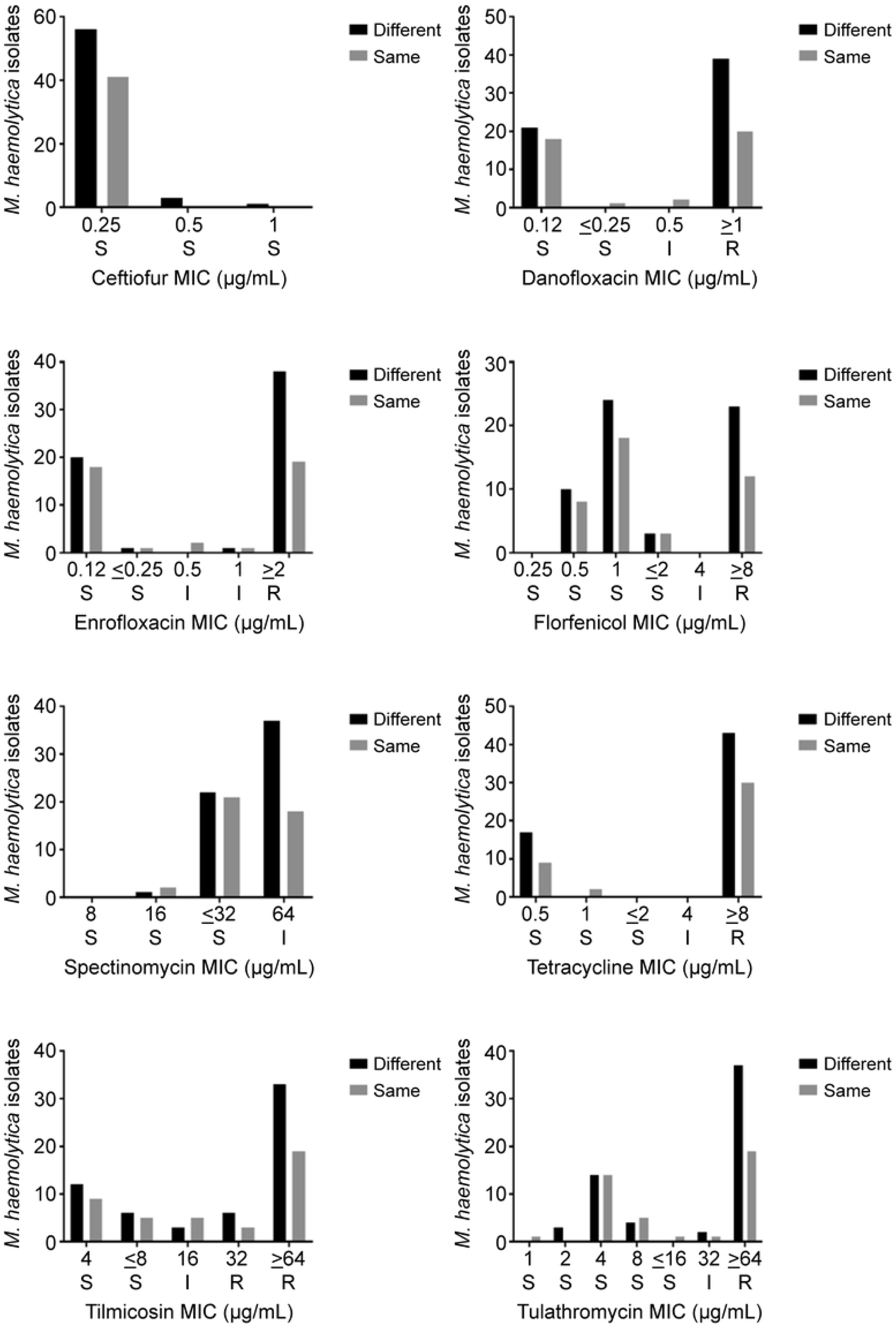
Distribution of minimum inhibitory concentrations (MIC) of antimicrobial agents with CLSI approved breakpoints for *M. haemolytica* in cattle receiving the same or different treatments. S = susceptible; I = intermediate; R = resistant. For “same” treatments, the first and second treatment were either bactericidal drugs or bacteriostatic drugs. For “different” treatments, the first treatment was bactericidal and second was bacteriostatic or *vice versa*.

Antimicrobial treatments were grouped based on their anticipated impact on bacterial growth in vitro, i.e., bactericidal (“cidal”) or bacteriostatic (“static”). We created a heatmap to illustrate the impact of specific pairs of combinations of first and second antimicrobial treatments on the number of isolates identified as resistant against the listed antimicrobials with CLSI breakpoints (Fig. 2). Red indicates the observed maximum number of resistant isolates and white (i.e., blank) represents no observation of antimicrobial resistance for a specific antimicrobial combination (Fig. 2). A similar examination of the data was not conducted for *P. multocida* and *H. somni* because there were an insufficient number of isolates for this to be meaningful.

**Fig. 2.**
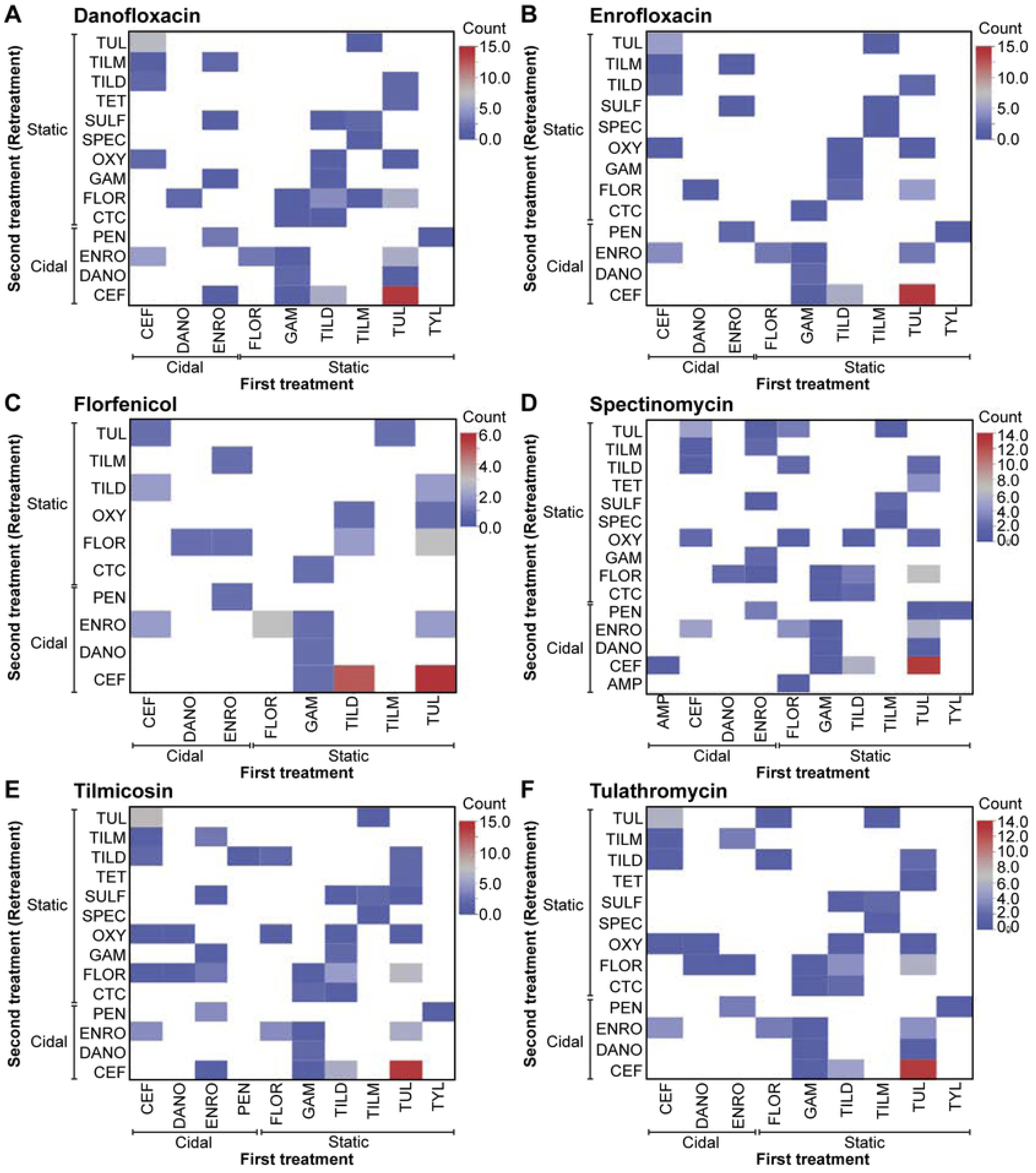
Heat maps showing pairwise interactions of antimicrobial treatment combinations associated with the isolation of resistant *M. haemolytica* organisms. The effect of treatment with ceftiofur (CEF), danofloxacin (DANO), enrofloxacin (ENRO), florfenicol (FLOR), gamithromycin (GAM), oxytetracycline (OXY), penicillin (PEN), spectinomycin (SPEC), sulfadimethoxine (SULF), tetracycline (TET), tildipirosin (TILD), tilmicosin (TILM), tulathromycin (TUL), and tylosin (TYL) as either first (X-axis) or second (Y-axis) treatment on the frequency of isolating *M. haemolytica* organisms resistant to danofloxacin (A), enrofloxacin (B), florfenicol (C), spectinomycin (D), tilmicosin (E) and tulathromycin (F) was examined using CLSI interpretive criteria. White indicates no observation of antimicrobial resistance with that specific combination.

Due to the limited number of isolates available from animals that received more than 2 treatments, we did not explore or conduct sensitivity analysis on the impact of other possible antimicrobial combinations on the isolation of resistant organisms. We also did not explore alternatives to the priors chosen for the Bayesian analysis, as we considered the chosen priors to be the most biologically defensible. A variable to account for the non-independence of isolates from the same animal was not included in the model, as the number of these cases was relatively small.

The distribution of AMR in bacterial isolates demonstrated an association between the isolation of an AMR bacteria and the number of treatments used (Fig. 3 and Table 3). The data indicate that administration of two or more antimicrobial agents to treat BRD in cattle may increase the likelihood of isolating an antimicrobial resistant pathogen (Fig. 3).

**Table 3.** 95% credible intervals (95% CIs) for the posterior distributions representing the probability of having at least one resistance result to at least one of the assessed antimicrobials (i.e., ρ) based on CLSI breakpoints stratified by the number of antimicrobials the animal received.

| Objective, Model | Percentile |  |  |
| --- | --- | --- | --- |
| Objective 1 Model 1: treatment frequency (n = 781) | 2.5% | 50% | 97.5% |
| 0 treatments | 0.29 | 0.36 | 0.55 |
| 1 treatment | 0.49 | 0.55 | 0.61 |
| 2 treatments | 0.63 | 0.69 | 0.76 |
| 3 treatments | 0.69 | 0.80 | 0.88 |
| 4+ treatments | 0.61 | 0.78 | 0.90 |
| Two treatment sequences (n = 211) |  |  |  |
| Same (bactericidal + bactericidal, bacteriostatic + bacteriostatic) | 0.61 | 0.71 | 0.79 |
| Different (bacteriostatic + bactericidal, bactericidal + bacteriostatic) | 0.59 | 0.69 | 0.76 |
| Four treatment sequences (n = 211) |  |  |  |
| Bactericidal + bactericidal | 0.54 | 0.77 | 0.92 |
| Bactericidal + bacteriostatic | 0.51 | 0.64 | 0.76 |
| Bacteriostatic + bacteriostatic | 0.59 | 0.69 | 0.79 |
| Bacteriostatic + bactericidal | 0.60 | 0.72 | 0.82 |

**Fig. 3.**
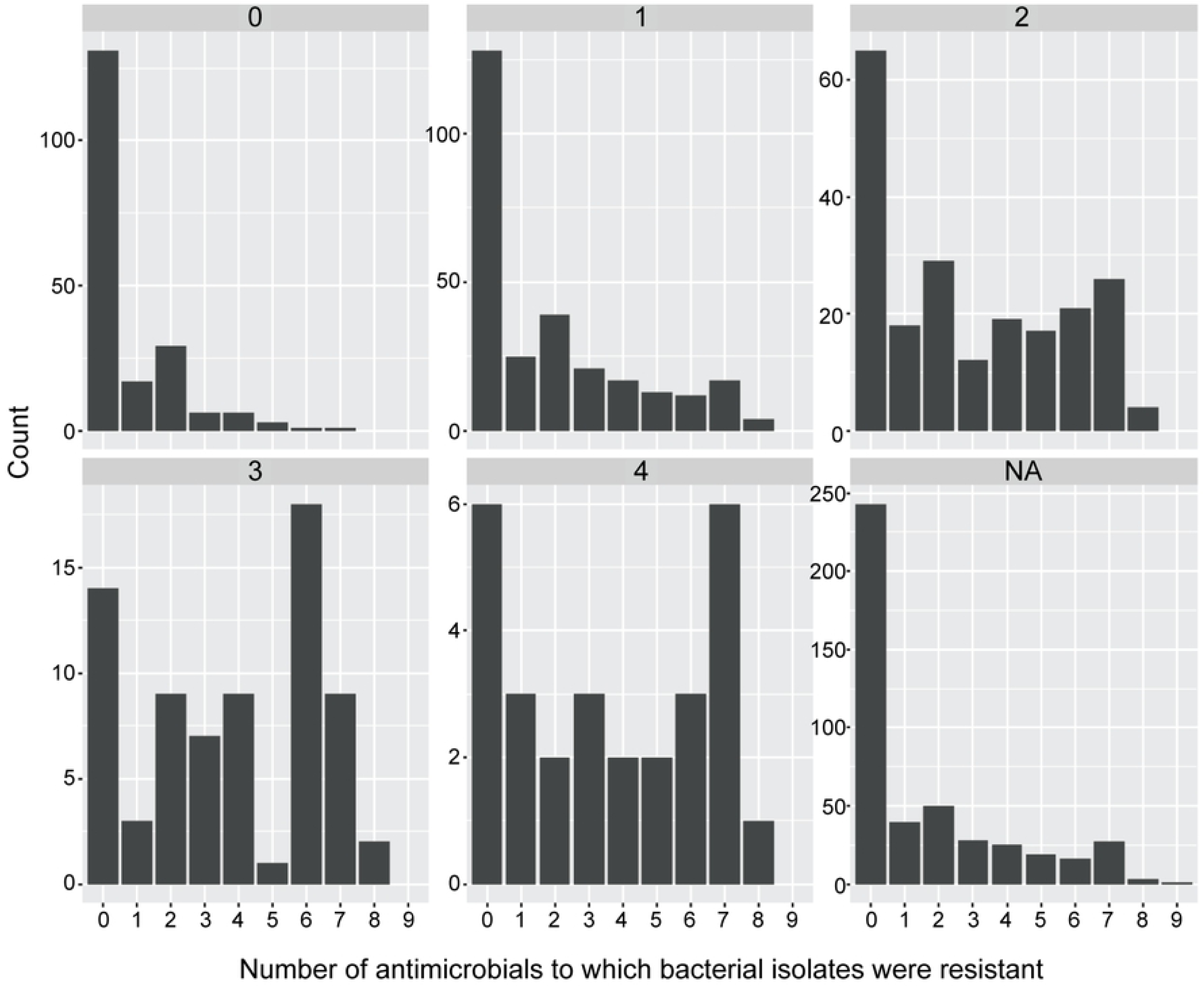
Observed frequency distribution of antimicrobial resistant isolates based on CLSI breakpoints for animals receiving antimicrobials for BRD. 0 = no treatment, 1 = 1 treatment, 2 = 2 treatments, 3 = 3 treatments, 4 = 4 or more treatments, and NA = missing information.

**Figure. 4.**
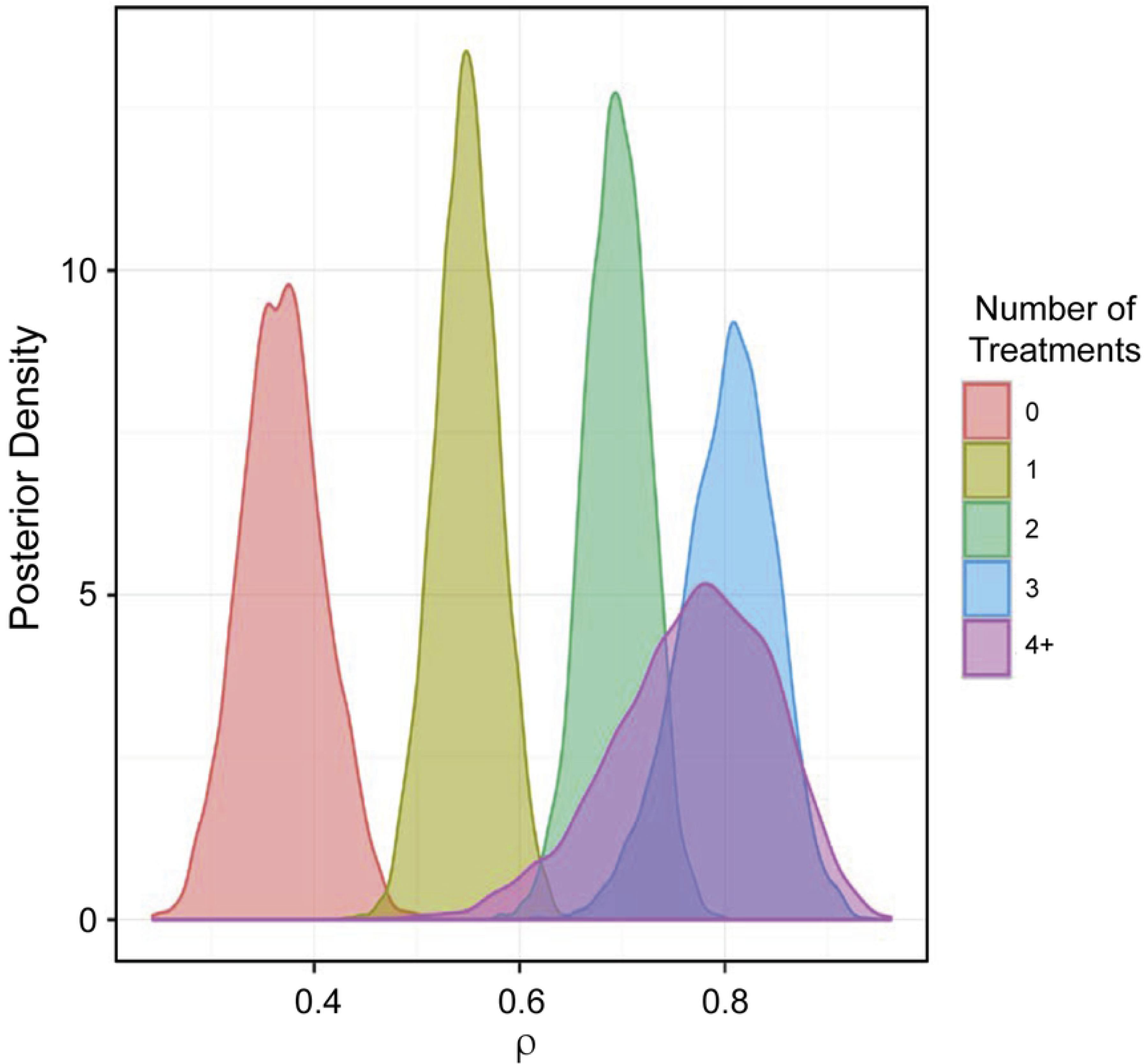

**Fig. 5.**
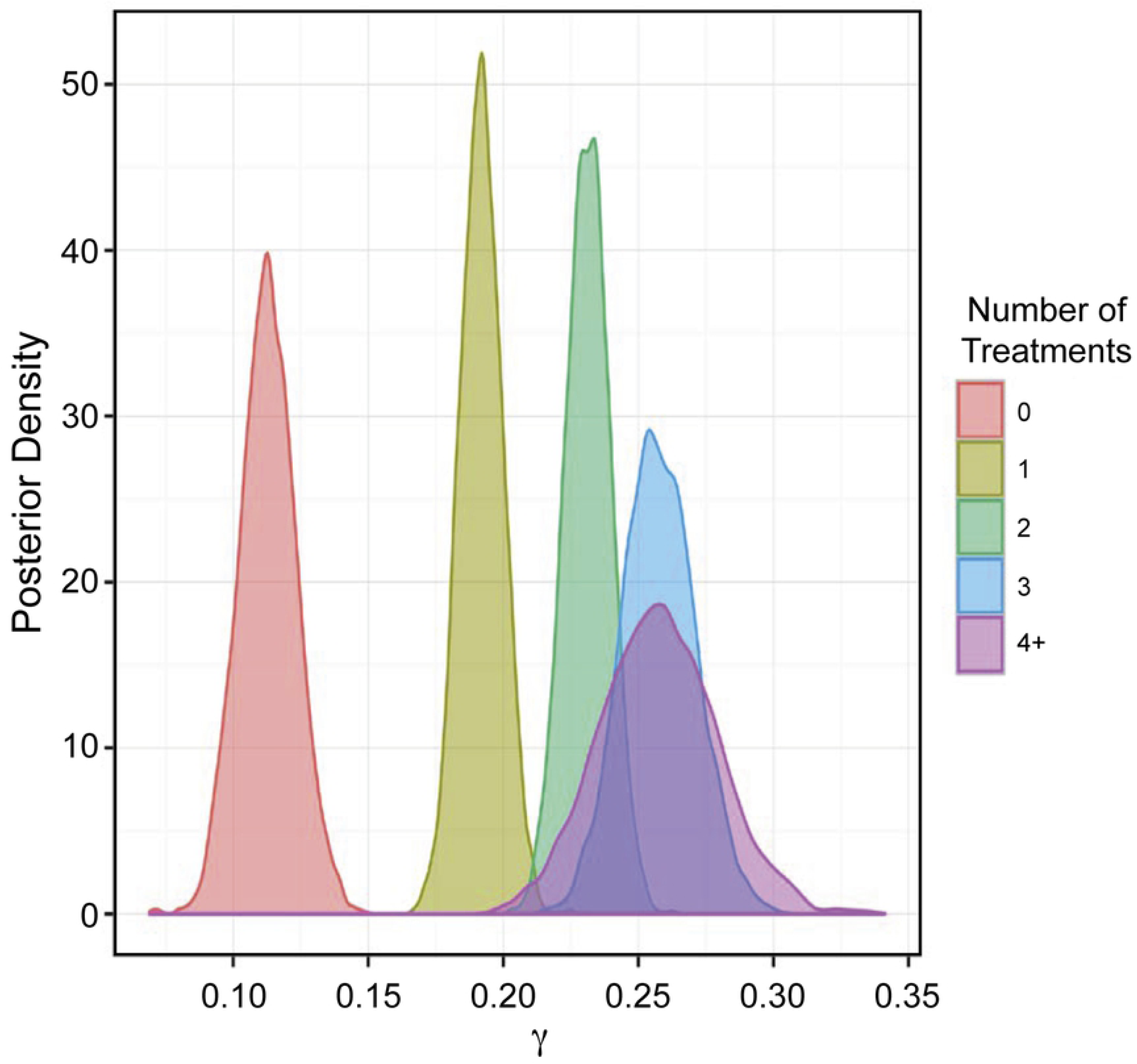
Posterior distributions of the probability that the isolate is resistant to multiple antimicrobials (i.e., γ_i_) stratified by treatment frequency. 0 = no treatment, 1 = 1 treatment, 2 = 2 treatments, 3 = 3 treatments, and 4+ = 4 or more treatments.

**Fig. 6.**
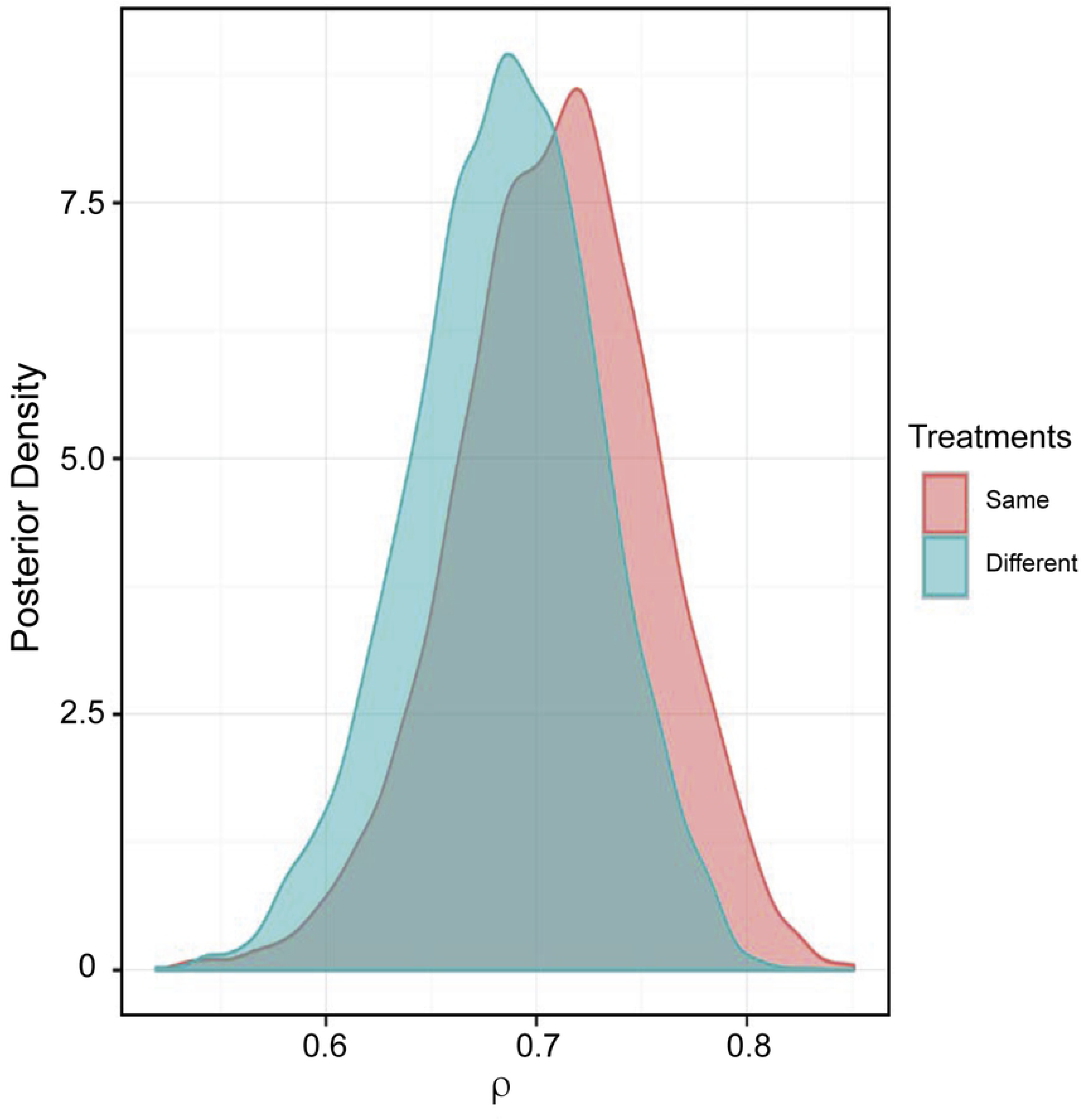
Posterior distribution of the probability that the isolate is resistant to at least one antimicrobial (i.e., ρ_i_) stratified by the expected in-vitro activity (i.e. bactericidal or bacteriostatic) of first and second treatment. Same = same in-vitro effect on bacterial growth, i.e., bactericidal followed by bactericidal or bacteriostatic followed by bacteriostatic; or different = different in-vitro effect on bacterial growth, i.e., bactericidal followed by bacteriostatic or bacteriostatic followed by bactericidal.

**Fig. 7.**
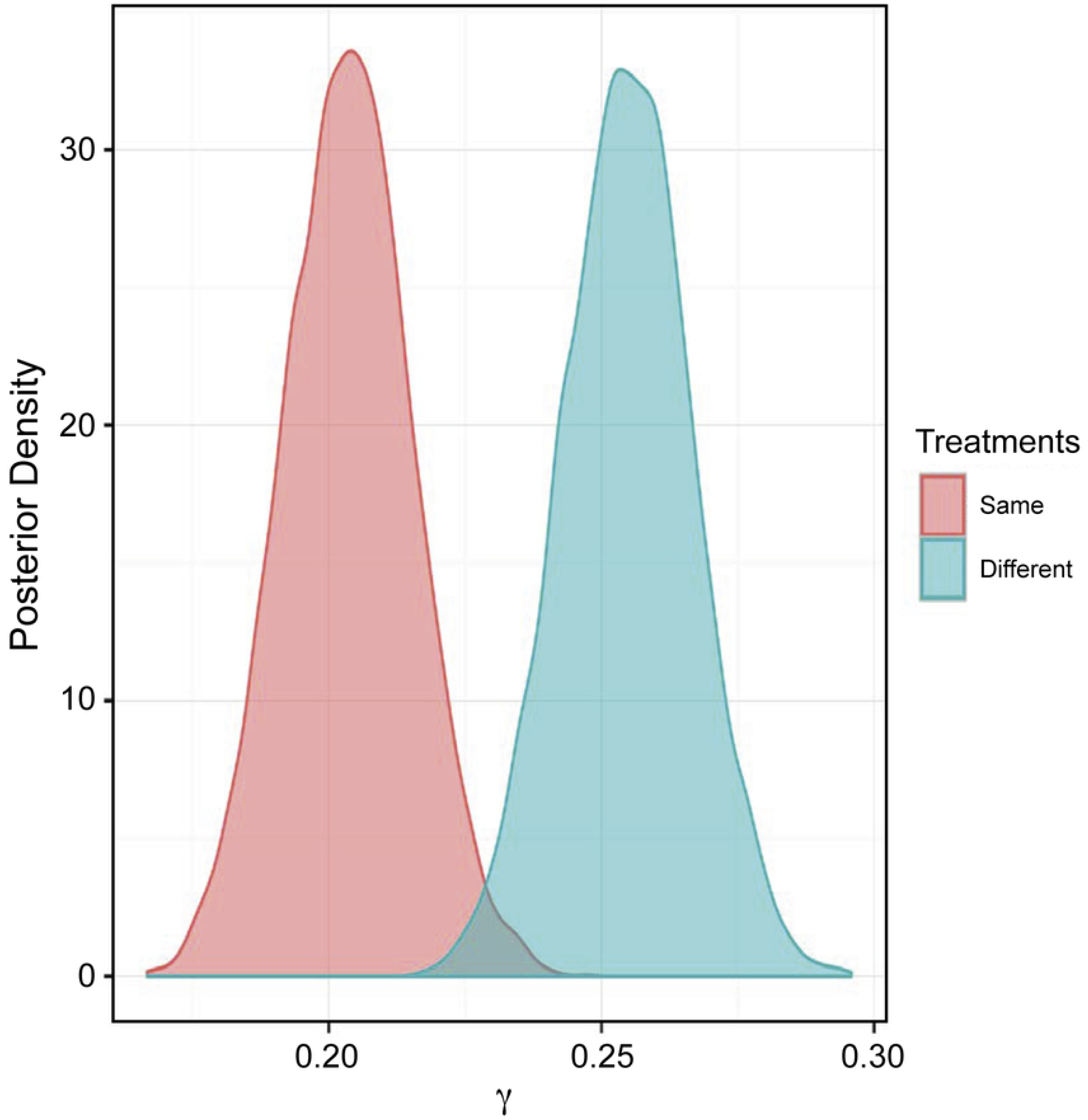
Posterior distributions of the probability that the isolate is resistant to multiple antimicrobials (i.e., γ_i_) stratified by the expected in-vitro activity (i.e. bactericidal or bacteriostatic) of first and second treatment. Same = same in-vitro effect on bacterial growth, i.e., bactericidal followed by bactericidal or bacteriostatic followed by bacteriostatic or different = different in-vitro effect on bacterial growth, i.e., bactericidal followed by bacteriostatic or bacteriostatic followed by bactericidal.

**Fig. 8.**
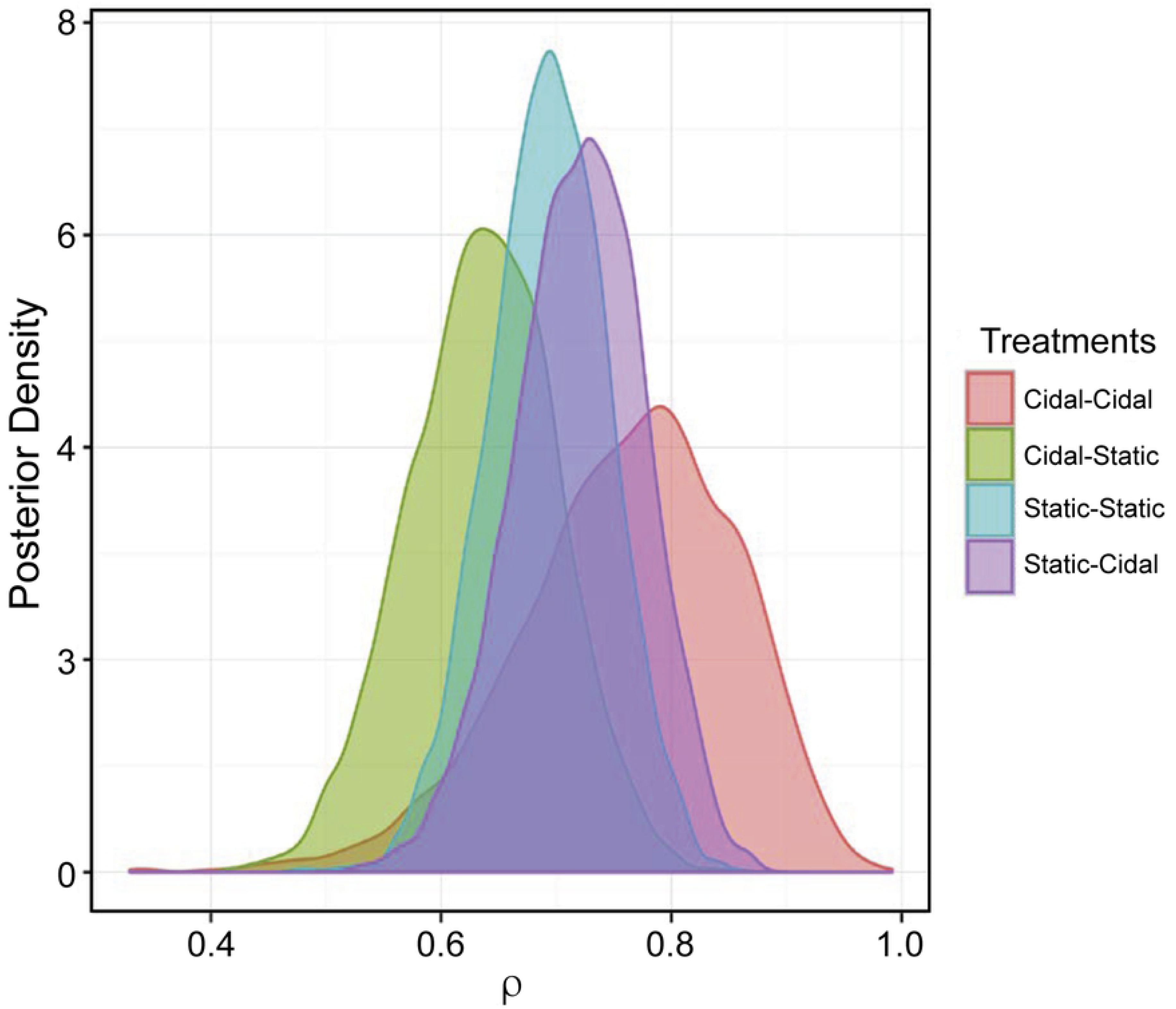
Posterior distribution of the probability that the isolate is resistant to at least one antimicrobial (i.e., ρ_i_) stratified by the expected in-vitro activity (i.e. bactericidal or bacteriostatic) of first and second treatment. Cidal-Cidal = bactericidal first treatment followed by bactericidal retreatment, Cidal-Static = bactericidal first treatment followed by bacteriostatic retreatment, Static-Static = bacteriostatic first treatment followed by bacteriostatic retreatment and Static-Cidal = bacteriostatic first treatment followed by bactericidal retreatment.

**Fig. 9.**
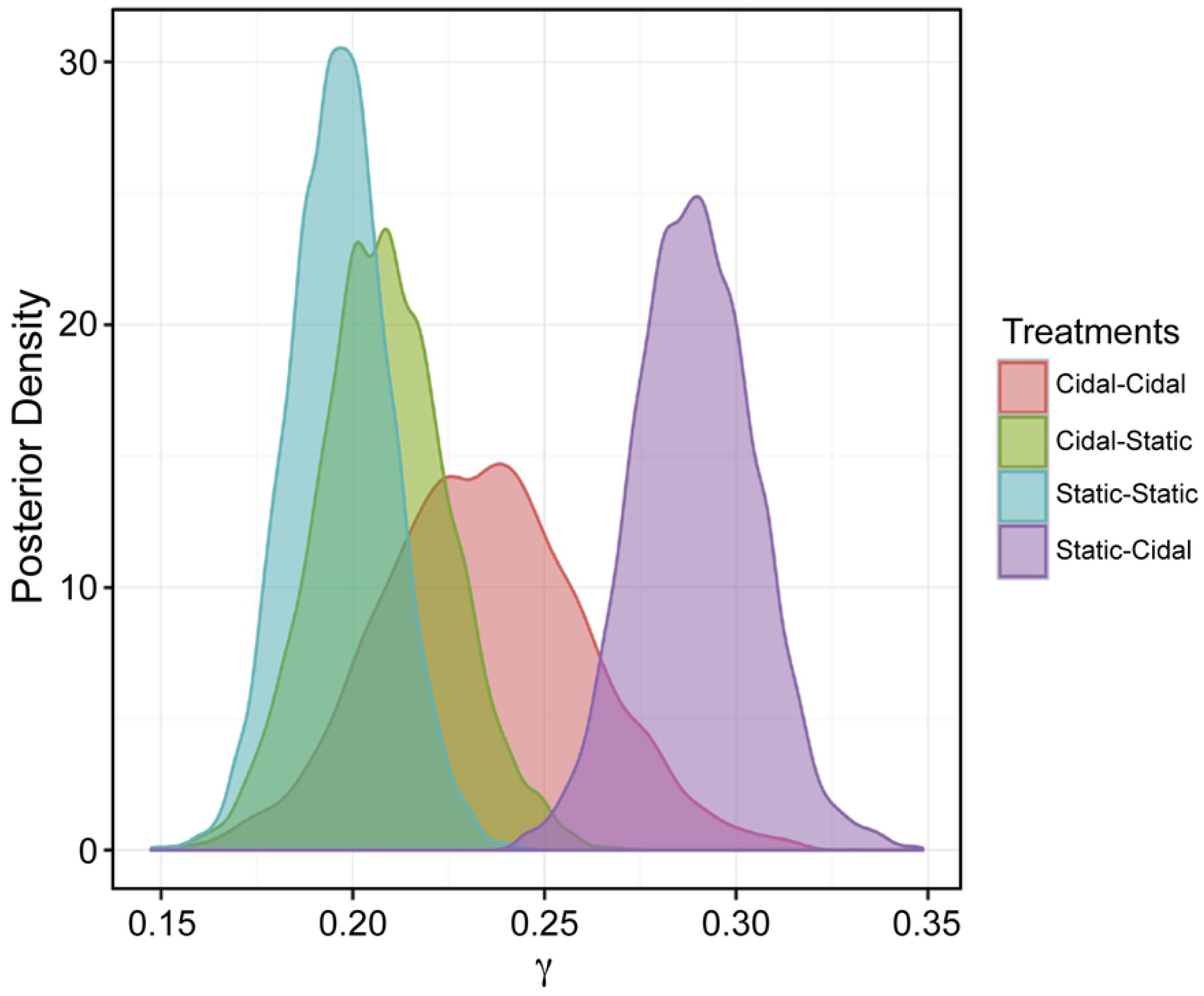
Posterior distributions of the probability that the isolate is resistant to multiple antimicrobials (i.e., γ_i_) stratified by the expected in-vitro activity (i.e. bactericidal or bacteriostatic) of first and second treatment. Cidal-Cidal = bactericidal first treatment followed by bactericidal retreatment, Cidal-Static = bactericidal first treatment followed by bacteriostatic retreatment, Static-Static = bacteriostatic first treatment followed by bacteriostatic retreatment and Static-Cidal = bacteriostatic first treatment followed by bactericidal retreatment.

For Objective 1, the posterior distribution for ρ (i.e., the probability of being resistant to at least one antimicrobial) is provided in **Error! Reference source not found.**The 95% CI for the ρ distribution is provided in **Error! Reference source not found. 3.** Based on the interpretation of the posterior distributions, the use of antimicrobials was associated with an increased probability of having at least one resistant outcome because the median and 95% CIs shift to the right toward higher probabilities as the number of antimicrobial treatments increased. In addition, there was evidence of an exposure response (i.e., increasing the number of treatments increases the probability of at least one resistant test). The evidence for a response to increasing antimicrobial exposure can be found in **Error! Reference source not found. 4**.

As reported in Table 4, ρ increased as the number of reported treatments increased. For example, 40%, 82%, 99%, and 100% of the time, ρ was higher if animals received more than 4 treatments when compared to ρ for 3 treatments, 2 treatments, 1 treatment, or 0 treatments, respectively. When ρ values were entered into the Bernoulli distribution, they translated into a higher prevalence of isolates with at least one resistant outcome.

**Table 4.** Posterior probability that the posterior distribution of *p_i+1_* > *p_i_*.

| Objective 1<br>Model 1 | 0 treatments | 1 treatment | 2 treatments | 3 treatments | 4+<br>treatments |
| --- | --- | --- | --- | --- | --- |
| 0 treatments | NA | 0.99 | 1 | 1 | 1 |
| 1 treatment | - | NA | 0.99 | 1 | 0.99 |
| 2 treatments | - |  | NA | 0.95 | 0.82 |
| 3 treatments | - | - |  | NA | 0.40 |
| 4+ treatments | - | - | - |  | NA |

$i$ represents the number of treatment approaches that differ based on Objective 1 Model 1.

The posterior distributions of γ (where γ_i_ = number of resistant tests each isolate has) are shown in **Error! Reference source not found.**. **5** (95% CI in Table 5).

Consistent with the results for ρ, there was evidence that increased exposure to antimicrobials resulted in a higher probability of an isolate being resistant to more than one antimicrobial (**Error! Reference source not found. 6).** However, for the difference between 3 treatments and 4+ treatments, there was only 49% probability (50/50) of one being higher than the other, suggesting a possible threshold or an imprecise estimate of the γ_i_ posterior distribution.

**Table 5.** Credibility percentiles for posterior distributions for the number of resistant test results from an isolate (γ_i_).

|  | Percentile |  |  |
| --- | --- | --- | --- |
| Objective 1 Model 1: Treatment frequency | 2.5% | 50% | 97.5% |
| 0 treatments | 0.09 | 0.11 | 0.13 |
| 1 treatment | 0.17 | 0.19 | 0.21 |
| 2 treatments | 0.21 | 0.23 | 0.25 |
| 3 treatments | 0.23 | 0.26 | 0.28 |
| 4+ treatments | 0.21 | 0.25 | 0.30 |
| Objective 2 Model 1: 2-treatment sequences |  |  |  |
| Same (bactericidal + bactericidal, bacteriostatic + bacteriostatic) | 0.18 | 0.20 | 0.23 |
| Different (bacteriostatic + bactericidal, bactericidal + bacteriostatic) | 0.23 | 0.25 | 0.28 |
| Objective 2 Model 2: 4-treatment sequences |  |  |  |
| Bactericidal + bactericidal, | 0.18 | 0.23 | 0.29 |
| Bactericidal + bacteriostatic, | 0.17 | 0.21 | 0.24 |
| Bacteriostatic + bacteriostatic, | 0.17 | 0.19 | 0.22 |
| Bacteriostatic + bactericidal | 0.26 | 0.28 | 0.32 |

**Table 6.** Posterior probability that γ_i+1_ is greater than γ_i_ where i is the number of treatment approaches which differ based on objective 1 model 1 and γ_i_ = number of resistant tests for an isolate.

| Objective 1 Model 1: Posterior distribution based on treatment frequency |  |  |  |  |  |
| --- | --- | --- | --- | --- | --- |
| i | 0 treatments | 1 treatments | 2 treatments | 3 treatments | 4+ treatments |
| 0 treatments | NA | 1 | 1 | 1 | 1 |
| 1 treatment | - | NA | 0.99 | 1 | 0.99 |
| 2 treatments | - |  | NA | 0.95 | 0.87 |
| 3 treatments | - | - |  | NA | 0.49 |
| 4+ treatments | - | - | - |  | NA |

$i$ is the number of treatments administered based on Objective 1 Model 1.

Objective 2 examined the development of resistance based on whether the antimicrobial selected for the initial treatment and retreatment would be expected to kill the bacteria in-vitro (i.e. bactericidal) or inhibit the replication of the bacteria in-vitro (i.e. bacteriostatic). As shown in **Error! Reference source not found.. 6** and **Error! Reference source not found. 3,** the posterior distribution of ρ (i.e., the probability of the isolate being resistant to at least one antimicrobial) when animals received drugs of the same or different mechanistic classes does not appear to be associated with different distributions.

However, when examining the posterior distribution of γ (where γ_i_ = number of resistant tests for an isolate), the posterior probability of γ_different_ > γ_same_ was 99% (**Error! Reference source not found.. 7 and Error! Reference source not found. 5**).

The results of the analysis from Objective 2 Model 1 suggest that the sequential administration of antimicrobial treatments with different effects on bacterial growth may be associated with higher numbers of resistant isolates and elevated MIC outcomes. Objective 2 Model 2 explores whether the sequence of bactericidal and bacteriostatic treatments has an impact on the probability of recovering a resistant BRD isolate. This analysis suggests that there is little impact of the treatment scheme sequence on the probability of identifying an isolate that is resistant to at least one antimicrobial (ρ). The specific posterior distributions and the 95% CI of ρ are shown in **Error! Reference source not found.. 8** and **Error! Reference source not found. 3.** Similarly, the posterior probability of an organism being resistant to at least one antimicrobial is presented in Table 7.

**Table 7.** Posterior probability that *p*_i+1_ is greater than p_i_ (*i*-4-level treatment mechanism sequence for objective 2 model 2).

|  | Bactericidal +<br>bactericidal | Bactericidal +<br>bacteriostatic | Bacteriostatic<br>+ bacteriostatic | Bacteriostatic<br>+ bactericidal |
| --- | --- | --- | --- | --- |
| Bactericidal + bactericidal | NA | 0.16 | 0.28 | 0.35 |
| Bactericidal + bacteriostatic | -- | NA | 0.73 | 0.81 |
| Bacteriostatic + bacteriostatic | - | - | NA | 0.62 |
| Bacteriostatic + bactericidal | - | - | - | NA |

$i$ is the 4-level treatment scheme sequence for Objective 2 Model 2.

**Table 8.**
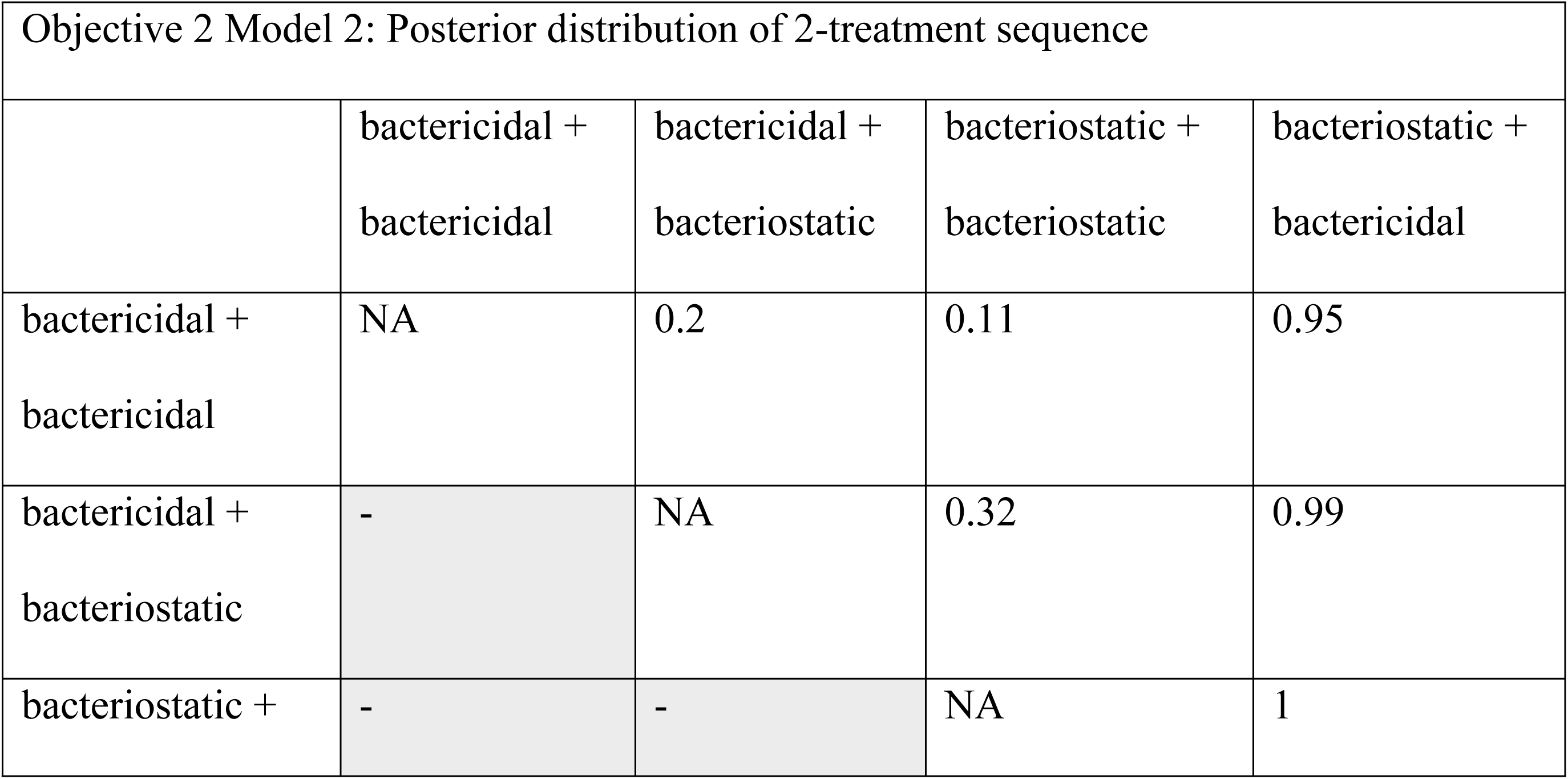
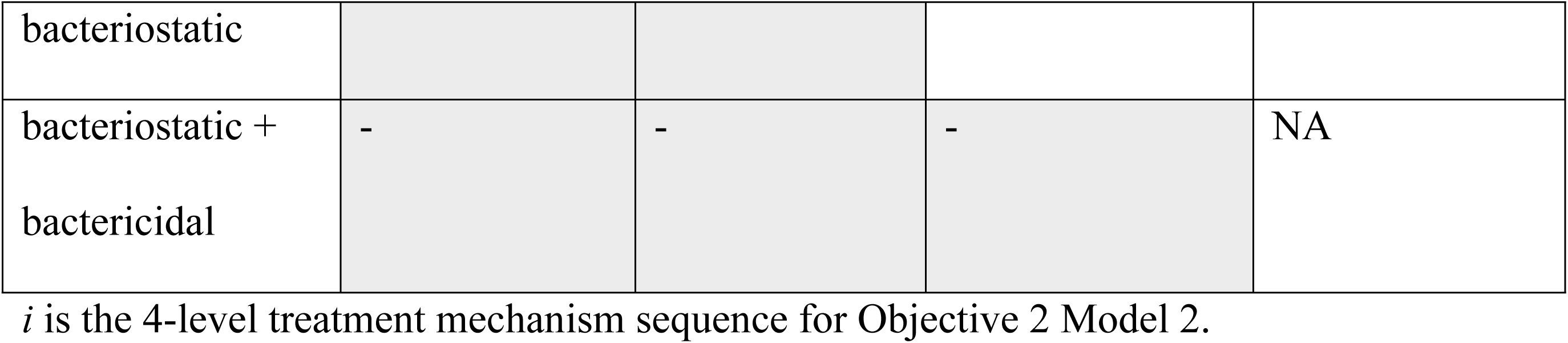
Posterior probability that the γ_i_ is greater γ_i-1_ (*i*-4-level treatment mechanism sequence for objective 2 model 2) where γ_i_ = number of resistant tests for an isolate.

As reported in Table 7, the probability of an organism being resistant to at least one antibiotic (ρ) was similar for the different treatment combinations. Specifically, in 62%, 81%, and 35% of cases, the probability of an organism being resistant to at least one antibiotic was higher if animals received a bacteriostatic antimicrobial for first treatment followed by a bactericidal antimicrobial for retreatment of BRD when compared to bacteriostatic-bacteriostatic, bactericidal-bacteriostatic, and bactericidal-bactericidal treatment, respectively.

With respect to the treatment, posterior gamma (γ) distributions shifted to the right in animals that received a first line, bacteriostatic antimicrobials followed by retreatment with a bactericidal antimicrobial (**Error! Reference source not found.. 9)**. This suggests that BRD pathogens isolated from these animals would be more likely to test resistant to more than one antimicrobial (Table 5). The probability of obtaining a resistant isolate from an animal receiving first-line bacteriostatic treatment followed by retreatment with a bactericidal antimicrobial being higher than the other sequences was >95% (**Error! Reference source not found. 8**).

These exploratory data suggest that treatment protocols stipulating first-line treatment with a bacteriostatic antimicrobial followed by retreatment with a bactericidal antimicrobial may be associated with an increased frequency of resistant BRD pathogen isolation. This observation may be due to the fact that bacteriostatic activity may antagonize the effect of bactericidal drugs. More specifically, bactericidal drugs act on bacteria that are in a growth phase; thus, the inhibitory activity of a bacteriostatic drug on the replication of bacteria may result in diminished activity of a subsequent bactericidal treatment [26].

To our knowledge, this survey is the first report that specifically considers AMR in livestock in the context of retreatments by different classes of antimicrobials (i.e., bacteriostatic and bactericidal) as well as different individual drugs (i.e., tulathromycin versus enrofloxacin). As such, this report provides insights into potential critical control points for antimicrobial stewardship in livestock production systems. For example, the heat map (Fig. 2) highlights how various antimicrobial combinations may influence changes in AMR profiles. Specifically, the combination of tulathromycin as the first-line treatment and ceftiofur as the second-line treatment increased the number of resistant isolates for all antimicrobial agents tested. In contrast, the use of ceftiofur as the first-line treatment and tulathromycin as the second-line treatment also led to an increase in the number of resistant isolates, but not to the same degree as when tulathromycin was the first antimicrobial used. The use of tildipirosin as the first-line treatment and ceftiofur as the second-line treatment also caused an increased number of isolates showing resistant phenotypes, but this increase was not as great as when tulathromycin was the first-line treatment. Tulathromycin and tildipirosin are in the same class of antimicrobials and have similar mechanisms of action. Bacterial pre-exposure to antimicrobials has been implicated as an important risk factor for AMR evolution during subsequent antimicrobial treatments [24,27,28]. Recently, the effect of sequential antimicrobial treatments on the development of antimicrobial resistance has been demonstrated for *Pseudomonas aeruginosa* and *Klebsiella pneumonia* in-vitro. In these laboratory studies the emergence of antimicrobial resistance also varied with the classes and concentrations of antimicrobials used for pre-exposure and sequential treatments [28, 29].

Macrolides, such as tulathromycin and tildipirosin, are appealing as first-line treatments for the control of BRD (metaphylaxis) in high risk cattle due to their efficacy and long residence times in plasma and tissues. Therefore, tulathromycin is one of the most frequently used antimicrobial drugs for metaphylaxis in the US [5]. However, metaphylaxis treatment with tulathromycin has been associated with a high prevalence of multidrug resistant *M. haemolytica* shedding in cattle [23, 30]. One explanation for the elevated antimicrobial resistance of *M. haemolytica* and *P. multocida* is the long elimination half-life of macrolides results in prolonged exposure to low concentrations of the bacteriostatic agent, which may be contributing to the development of AMR. During minimal inter-treatment intervals of 3–5 days, macrolides are still present at appreciable concentrations that may allow for drug-drug interactions when a second antimicrobial treatment is administered. Therefore, longer inter-treatment intervals are recommended when using antimicrobials with longer elimination half-lives [17]. Unfortunately the interval between treatments was not recorded on the submission histories analyzed for the present study and so the impact of post-treatment interval on the emergence of AMR could not be assessed.

Our data suggest that the number of treatments as well as altering antimicrobial classes may impact antimicrobial resistance patterns. Damas *et al.* used three antimicrobial classes (penicillins, cephalosporins, and fluoroquinolones) to treat serious infections in intensive care unit patients for 8-month periods over a 2-year duration. After studying the effect of the sequential use of these three antimicrobial classes, antimicrobial rotation was associated with a higher risk for the development of antimicrobial resistance [31].

Differences between resistance patterns were displayed by the MIC distributions of *M. haemolytica*. With the exception of ceftiofur and florfenicol, the numbers of resistant *M. haemolytica* isolates were greater when different antimicrobial classes were used (**Figs. 1 & 2**). However, the number of susceptible *M. haemolytica* isolates did not differ with antimicrobial classes. In general, resistance to ceftiofur was rare, even in isolates obtained from animals treated with different antimicrobial classes. However, it is speculated that CLSI-approved breakpoints may not be accurate for ceftiofur against *M. haemolytica* and *P. multocida* for treatment of respiratory disease. It is known that exposure to antimicrobials offers an advantage to resistant mutants in competition with the susceptible wild-type population; however, the impact of multidrug combinations of different classes on positive selection of resistant mutants has not been closely examined [32]. In a previous report, authors measured the ratio change of doxycycline-resistant and doxycycline-sensitive *E. coli* following treatment with doxycycline alone or in combination with erythromycin. The doxycycline-resistant mutants outnumbered the susceptible wild-type population of *E.coli* in both treatment conditions, but there was greater selection for the resistant mutants with the combination treatment [33]. In line with these reports, our study also suggests that using a combination of different classes of antimicrobials may increase the risk of selection of resistant mutants.

Our ability to assess the impact of different drug classes on AMR for *P. multocida* and *H. somni* was limited in the present study due to the relatively small number of isolates with associated treatment histories that were available for analysis. However, it is known that the MIC distribution for *P. multocida* and *H. somni* may not have the same pattern as *M. haemolytica* isolates. In a previous report, pre-exposure to tulathromycin developed bacterial resistance in *M. haemolytica* but not in *P. multocida* [34]. The number of *M. haemolytica* isolates compared to the number of *P. multocida* and *H. somni* isolates may influence the observations of this study. Regardless, the use of different mechanistic classes of antimicrobials may lead to a greater number of resistant isolates. Van Loon *et al*. reported that bacteria exhibit reduced susceptibility during treatment with variation in the classes of antimicrobials [35].

Though drug resistance has been a concern of scientists for decades, and specific BRD pathogen resistance was first reported over 40 years ago [36], our study appears to be the first thorough investigation of the effects of treatment number and type on subsequent AMR isolates in cattle with BRD. More recent investigations of resistance to individual drugs are more extensive and characterize AMR among BRD pathogens after treatment with tetracyclines, macrolides, beta-lactams, fluoroquinolones, sulphonamides, phenicols, aminoglycosides, and lincosamides [8,11,12,13,14]. The current study is representative, though non-exhaustive, and exhibits a novel approach for AMR microbe analysis in BRD because multiple treatments of various drugs and antimicrobial classes are evaluated.

The impact of multiple antimicrobial treatments represents an understudied area of research in veterinary medicine. It has been shown that feedlot cattle are routinely treated with antimicrobials more than once if the initial response is inadequate; however, cattle that receive multiple antimicrobial treatments exhibit higher mortality rates from disease [5]. Furthermore, animals that fail to respond to the initial treatment with one class of drug (e.g., bacteriostatic) are usually retreated with a different class of drug (e.g., bactericidal), which suggests a lack of consensus on any particular retreatment protocol [5]. This lack of consensus is likely due to the scarcity of literature on pathogen response to multiple treatment regimens with different classes of antimicrobial agents. Our study suggests that sequential treatment with different classes of antimicrobials is a risk factor for developing drug resistance. Therefore, a review of antimicrobial pre-exposure should be taken before the initiation of subsequent antimicrobial therapy to prevent the emergence of antimicrobial resistance in cattle infected with BRD.

As concern about the impact of AMR microbes on animal and public health increase, additional knowledge from studies such as the current one are needed to investigate interventions that reduce the development of antimicrobial resistance. Furthermore, a microbiological diagnosis should be established before using broad-spectrum antimicrobials to treat BRD of unknown etiology. Unfortunately, the amount of time it takes to obtain AMR isolate results and the associated costs are two major limitations for the use of laboratory microbiology in veterinary medicine [25]. Furthermore, this study demonstrates the value and importance of including comprehensive treatment histories to accompany the submission of veterinary diagnostic laboratory samples. The current study of antimicrobial sensitivity patterns in a region can guide veterinarians to choose safer and more effective treatment protocols. Future studies on antimicrobial resistance could facilitate decision-making when animals contained in feedlots exhibit chronic illness and there is the potential need for multiple treatments with antimicrobial agents.

### Key results

These exploratory data suggest that treatment protocols stipulating first-line treatment with a bacteriostatic followed by second-line treatment with a bactericidal may increase the probability that drug resistance develops. As concern about antimicrobial resistance increases from an animal and public health perspective, this knowledge suggests potential ways to reduce the development of resistance. The hypothesis that the impact of an antimicrobial on bacterial growth may be associated with the risk of increased resistance needs to be tested in a clinical trial. Such a trial would also need to determine whether treatment efficacy is affected by a change in treatment protocol or post-treatment interval. If treatment effectiveness proves to be the same, then we may have an avenue by which to reduce the induction of resistance via the recommendation that veterinarians tailor their treatment regimens to reduce the potential for AMR development.

### Strengths and limitations

Although this study is hypothesis-generating, it has several strengths. The data set is reasonably large for the questions we asked. Although a great deal of data were missing, we limited our analysis to specific questions to avoid impact due to this missing data. Furthermore, we recognized the limits of the passively collected and hypothesis-generating nature of the data by not formally testing a hypothesis. The zero-inflated beta-binomial model that we used is an intuitive model that fit the underlying data well. We could not adjust this model for any confounders because of missing data; however, given the cross-sectional nature of the data, any attempt to adjust for confounders to improve causal inference would have been misleading and was thus avoided.

### Interpretation and generalizability

Our overall interpretation of the data suggests that there is direct correlation between the number of treatments to which an animal was exposed and the emergence of treatment resistance. In addition, sequential treatments of BRD and the use of antimicrobials with different mechanisms of antibacterial activity (i.e., -static versus -cidal) may serve as a risk factor for the development of AMR.

## Authorship declarations

JFC: Conceived the study, provided study guidance and relevant interventions, interpreted the results, and prepared and approved the final manuscript.

DRM: Conceived the original retrospective work, participated in data collection and interpretation of the results, and approved the final manuscript.

LF: Conducted the Bayesian analysis, including writing the code.

PKS: Participated in compiling and interpreting the results and preparation of the manuscript, and approved the final manuscript.

AMS: Participated in compiling and interpreting the results and preparation of the manuscript, and approved the final manuscript.

ACK: Participated in the antimicrobial susceptibility testing, interpretation of the results, and preparation of the manuscript, and approved the final manuscript.

VLC: Participated in compiling and interpreting the results and preparation of the manuscript, and approved the final manuscript.

TJE: Participated in compiling and interpreting the results and preparation of the manuscript, and approved the final manuscript.

AOC: Conducted the descriptive analysis and assisted with the Bayesian analysis, prepared the draft of the statistical methods and results, and approved the final manuscript.

## Publication declaration

The authors declare that this is a full and accurate description of the project and no important information or analyses are omitted. A second paper provides a detailed description of the results for the individual antimicrobials

## Acknowledgments

We would like to thank Pete Walsh for helping with some of the descriptive coding.

## Sources and manufacturers

a. BOPO6F, Thermo Scientific, Oakwood Village, OH.
b. Sensititre AIM, Thermo Scientific, Oakwood Village, OH.
c. Sensititre Vizion System, Thermo Scientific, Oakwood Village, OH.

## Data availability statement

The data that support the findings of this study are available from the Iowa State University Veterinary Diagnostic Laboratory. Restrictions apply to the availability of these data, which are not publicly available due to client confidentiality. Data are available from the authors with the permission of Iowa State University.

## References

1. Griffin D Economic impact associated with respiratory disease in beef cattle. Vet Clin North Am Food Anim Pract. 1997; 13: 367–77.

2. Snowder GD, Van Vleck LD, Cundiff LV, Bennett GL, Koohmaraie M, Dikeman ME. Bovine respiratory disease in feedlot cattle: Phenotypic, environmental, and genetic correlations with growth, carcass, and longissimus muscle palatability traits. J. Anim. Sci. 2007; 85: 1885–1892.

3. Sanderson MW, Dargatz DA, Wagner BA Risk factors for initial respiratory disease in United States’ feedlots based on producer-collected daily morbidity counts. Can Vet J. 2008; 49: 373–8.

4. Miles DG, Rogers KC BRD control: tying it all together to deliver value to the industry. Anim Health Res Rev. 2014;15: 186–8.

5. USDA–APHIS–VS: National Animal Health Monitoring System Beef Feedlot Study 2011. Part IV: Health and Health Management on U.S. Feedlots with a Capacity of 1,000 or More Head. 2013. Available from: https://www.aphis.usda.gov/animal_health/nahms/feedlot/downloads/feedlot2011/Feed11_dr_PartIV.pdf

6. Fales WH, Selby LA, Webber JJ, Hoffman LJ, Kintner LD, Nelson SL, et al. Antimicrobial resistance among Pasteurella spp recovered from Missouri and Iowa cattle with bovine respiratory disease complex. J Am Vet Med Assoc.1982; 181:477–9.

7. Welsh RD, Dye LB, Payton ME, Confer AW Isolation and antimicrobial susceptibilities of bacterial pathogens from bovine pneumonia: 1994--2002. J Vet Diagn Invest. 2004; 16: 426–31, 2004.

8. Klima CL, Alexander TW, Read RR, Gow SP, Booker CW, Hannon S, Selinger LB Genetic characterization and antimicrobial susceptibility of Mannheimia haemolytica isolated from the nasopharynx of feedlot cattle. Vet Microbiol. 2011; 149(3-4): 390–398.

9. Lubbers BV, Hanzlicek GA Antimicrobial multidrug resistance and coresistance patterns of Mannheimia haemolytica isolated from bovine respiratory disease cases--a three-year (2009-2011) retrospective analysis. J Vet Diagn Invest. 2013; 25: 413–7.

10. Magstadt DR, Schuler AM, Coetzee JF, Krull AC, O’Connor AM, Cooper VL, Engelken TJ. Treatment history and antimicrobial susceptibility results for Mannheimia haemolytica, Pasteurella multocida, and Histophilus somni isolates from bovine respiratory disease cases submitted to the Iowa State University Veterinary Diagnostic Laboratory from 2013-2015. J Vet Diagn Invest. 2018; 30: 99–104.

11. Watts JL, Sweeney MT Antimicrobial resistance in bovine respiratory disease pathogens: measures, trends, and impact on efficacy. Vet Clin North Am Food Anim Pract. 2010; 26: 79–88.

12. Michael GB, Kadlec K, Sweeney MT, Brzuszkiewicz E, Liesegang H, Daniel R, et al. ICE Pmu1, an integrative conjugative element (ICE) of Pasteurella multocida: analysis of the regions that comprise 12 antimicrobial resistance genes. J Antimicrob Chemother. 2011; 67(1): 84–90.

13. Pardon B, Hostens M, Duchateau L, Dewulf J, De Bleecker K, Deprez P. Impact of respiratory disease, diarrhea, otitis and arthritis on mortality and carcass traits in white veal calves. BMC Vet Res. 2013; 9(1): 79.

14. Woolums AR, Karish BB, Frye JG, Epperson W, Smith DR, Blanton Jr, et al. Multidrug resistant *Mannheimia haemolytica* isolated from high-risk beef stocker cattle after antimicrobial metaphylaxis and treatment for bovine respiratory disease. Vet Microbiol. 2018; 221:143–152.

15. Clinical and Laboratory Standards Institute (CLSI). Performance standards for antimicrobial disk dilution susceptibility tests for bacteria isolated from animals. CLSI, Wayne, PA16; 2008.

16. Johnson KK and Pendell DL. Market Impacts of Reducing the Prevalence of Bovine Respiratory Disease in United States Beef Cattle Feedlots. Front Vet Sci. 2017; 4: 189. doi: 10.3389/fvets.2017.00189

17. Apley, MD. Treatment of calves with bovine respiratory disease: duration of therapy and posttreatment intervals. Vet Clin North Am Food Anim Pract. 2015; 31: 441–453.

18. Anholt RM, Klima C, Allan N, Matheson-Bird H, Schatz C, Ajitkumar P, Otto SJG, Peters D, Schmid K, Olson M, McAllister T and Ralston B. Antimicrobial Susceptibility of Bacteria That Cause Bovine Respiratory Disease Complex in Alberta, Canada. Front Vet Sci. 2017; 4: 207. doi: 10.3389/fvets.2017.00207

19. Portis E, Lindeman C, Johansen L, Stoltman G. A ten-year (2000–2009) study of antimicrobial susceptibility of bacteria that cause bovine respiratory disease complex— *Mannheimia haemolytica, Pasteurella multocida*, and *Histophilus somni*—in the United States and Canada. J Vet Diagn Invest. 2012; 24: 932–44. doi:10.1177/1040638712457559

20. Klima CL, Zaheer R, Cook SR, Booker CW, Hendrick S, Alexander TW, et al. Pathogens of bovine respiratory disease in North American feedlots conferring multidrug resistance via integrative conjugative elements. J Clin Microbiol. 2014; 52: 438–48. doi:10.1128/JCM.02485-13

21. Noyes N, Benedict K, Gow S, Booker C, Hannon S, McAllister T, et al. *Mannheimia haemolytica* in feedlot cattle: prevalence of recovery and associations with antimicrobial use, resistance, and health outcomes. J Vet Intern Med. 2015; 29:705–13. doi:10.1111/jvim.12547

22. DeDonder KD, Apley MD. A literature review of antimicrobial resistance in pathogens associated with bovine respiratory disease. Anim Health Res Rev. 2015; 16:125–34.

23. Snyder E., Credille B., R. Berghaus, Giguere S. Prevalence of multi drug antimicrobial resistance in isolated from high-risk stocker cattle at arrival and two weeks after processing. J Anim Sci. 2017; 95: 1124–1131.

24. Gould IM, MacKenzie FM. Antibiotic exposure as a risk factor for emergence of resistance: the influence of concentration. J Appl Microbiol. 2002;92: 78S–84S.

25. De Briyne N, Atkinson J, Pokludová L, Borriello SP, Price S. Factors influencing antibiotic prescribing habits and use of sensitivity testing amongst veterinarians in Europe. Vet Rec. 2013; 173: 475 http://dx.doi.org/10.1136/vr.101454

26. Ocampo PS, Lázár V, Papp B, Arnoldini M, Abel zur Wiesch P, Busa-Fekete R, Fekete G, Pál C, Ackermann M, Bonhoeffer S. Antagonism between bacteriostatic and bactericidal antibiotics is prevalent. Antimicrob Agents Chemother. 2014; 58(8): 4573–82. doi: 10.1128/AAC.02463-14.

27. var der Horst MA, Schuurmans JM, Smid MC, Koenders BB, ter Kuile BH. De vovo acquisition of resistance to three antibiotics by Escherichia coli. Microb Drug Resist. 2011;17(2): 141–7.

28. Vestergaard M, Paulander W, Marvig RL, Clasen J, Jochumsen N, Molin S, et al. Antibiotic combination therapy can select for broad-spectrum multidrug resistance in *Pseudomonas aeruginosa*. Int J Antimicrob Agent. 2016; 47(1): 48–55.

29. Kim J, Jo A, Chukeatirote E, Ahn J. Assessment of antibiotic resistance in *Klebsiella pneumoniae* exposed to sequential in vitro antibiotic treatments. Ann Clin Microbiol Antimicrob. 2016; 15:60. https://doi.org/10.1186/s12941-016-0173-x

30. Crosby S, Credille B, Giguère S, Berghaus R. Comparative efficacy of enrofloxacin to that of tulathromycin for the control of bovine respiratory disease and prevalence of antimicrobial resistance in *Mannheimia haemolytica* in calves at high risk of developing bovine respiratory disease. J Anim Sci. 2018; 96: 1259–1267. https://doi.org/10.1093/jas/sky054

31. Damas P, Canivet JL, Ledoux D, Monchi M, Melin P, Nys M, De Mol P. Selection of Resistance during sequential use of preferential antibiotic class. Intensive Care Med. 2006; 32; 67– 74.

32. Levy, S. B. & Marshall, B. Antibacterial resistance worldwide: causes, challenges and responses. Nature Med. 2004; 10, S122–S129.

33. Chait R, Craney A, Kishony R. Antibiotic interactions that select against resistance. Nature. 2007; 446(7136): 668–71. DOI: 10.1038/nature05685

34. Rajamanickam K, Yang J and Sakharkar MK. Gallic Acid Potentiates the Antimicrobial Activity of Tulathromycin Against Two Key Bovine Respiratory Disease (BRD) Causing-Pathogens. Front Pharmacol. 2019; 9: 1486. doi: 10.3389/fphar.2018.01486

35. Loon HJ van, Vriens MR, Fluit AC, Troelstra A, van der Werken C, Verhoef J, Bonten M. Antibiotic rotation and development of gram-negative antibiotic resistance. Am J Respir Crit Care Med. 2005; 171: 480–487.

36. Chang WH, Carter GR. Multiple drug resistance in Pasteurella multocida and Pasteurella haemolytica from cattle and swine. J Am Vet Med Assoc. 1976; 169(7):710–712.

